# Biomanufacturability of Squid Ring Teeth Protein Library via Orthogonal High-Throughput Screening

**DOI:** 10.1101/2025.08.25.672143

**Authors:** Khushank Singhal, Benjamin D. Allen, Thomas M. Baer, Melik C. Demirel

## Abstract

Structural proteins are valued by materials engineers for their mechanical properties, chemical functionalities, and biodegradability. Several studies focus on high-throughput screening of structural proteins, which is crucial for understanding and improving proteins by analyzing entire sequence spaces, as demonstrated in directed evolution. A major challenge is the poor biomanufacturability of recombinant structural proteins, often due to toxicity issues (e.g., aggregation, cell stress, inclusion bodies) limiting yield. High-throughput screening can help solve these issues and improve the biomanufacturability of structural proteins. Based on naturally observed squid ring teeth proteins, we introduce a structural protein library, enabling us to explore a broad sequence space. We selected 33 amino acid fragments with four repeats from six different native squid species, creating a recombinant protein library of about 1.2 million variants (i.e., 33^4^). We demonstrated an orthogonal screening method that combines fluorescent-assisted cell sorting (FACS) and fluorescent microcapillary-array-based screening to establish correlations from genotype to single cells and ensembles. This optical high-throughput approach enables screening across the spectrum from individual cells to groups, facilitating the enrichment of desired clones. Our workflow distinguishes between expression and growth traits, supporting systematic genetic design studies focused on biomanufacturability. We observed that brighter clones tend to contain self-similar sequences (i.e., perfect repeats of fragments), whereas dimmer clones in the library have less similarity (i.e., imperfect repeats). The ability to screen large structural protein libraries not only accelerates research and development but also creates new opportunities for materials research and advanced biotechnological processes, underscoring its importance in modern synthetic biology.

## INTRODUCTION

Structural proteins, such as silk,^1^ collagen,^2^ elastin,^3^ keratin,^4^ and squid ring teeth (SRT)^5^, have been utilized by materials engineers due to their exceptional mechanical properties, chemical functionalities, and biodegradability. Sequence-structure-property relationships are fundamental to protein-based materials, particularly those derived from structural proteins.^6^ Amino acid sequence influences protein folding, which in turn affects how the protein responds to environmental factors.^7^ Many structural features, such as hydrophobicity caused by exposed amino acids, crystalline or amorphous regions, protein entanglements, and charged functional groups, contribute to the unique properties of protein materials. The resulting properties of these proteins and their composites with inorganic materials^8^ are used in various fields, from materials science to biomedicine engineering.^9^

SRT has gained attention recently due to its unique composite structure and remarkable toughness, which enable quick and agile movements.^10^ SRT is used in defense and hunting strategies, where squid has developed various features that make them effective predators, such as strong tentacles and many SRTs along these tentacles.^11^ SRT proteins are multifunctional, exhibiting unique properties such as thermal conductivity switching,^12^ rapid self-healing,^13^ tunable hydrophobicity,^14^ enhanced optical properties,^15^ and triboelectric properties.^16^ This variability presents a valuable opportunity to optimize key factors affecting biomanufacturability, enabling the identification of sequence designs that enhance protein yield, solubility, and stability for large-scale production.

While SRT proteins and, in general, structural proteins^17–19^ have been recombinantly engineered and produced at a laboratory scale, many challenges still exist in their biomanufacturing.^20^ For example, the diversity of SRT protein sequences and their impact on expression and solubility are difficult to predict.^21^ Efficient methods are needed to screen and select structural protein variants with high expression and solubility, as traditional protein engineering techniques often have low throughput and rely on trial-and-error approaches. Arakawa et al. emphasized the importance of global sampling by cataloging silk gene sequences from 1098 spider species and analyzing how amino acid motifs influence properties such as mechanical strength, thermal stability, and hydration in over 446 dragline silks.^22^ The aforementioned study took several years to complete. Efficient protocols for protein library synthesis, like combinatorial methods, could supplement such studies by providing faster sequence-space coverage than previously thought. Rational design of structural proteins, inspired by natural protein sequences, enables further optimization, such as increasing the molecular weight of engineered proteins through intein-mediated polymerization.^23, 24^ However, these methods have been limited to a few protein sequences found in nature or their modifications, leaving room for significant improvements in their properties through modern synthetic biology techniques, especially by exploring large protein libraries.

Several studies have concentrated on high-throughput screening of proteins.^25–27^ Analyzing the entire sequence space for protein libraries is essential for understanding and improving proteins, as shown by directed evolution studies.^28–31^ Mutagenesis is a powerful technique in protein engineering that enables the modification of proteins to create libraries of variants.^32^ Site-directed mutagenesis enables precise changes at specific nucleotide positions to modify amino acids, a technique commonly used in rational protein design based on structural insights. Random mutagenesis, achieved through error-prone PCR or chemical mutagenesis, introduces a diverse range of mutations for screening. DNA shuffling is a directed evolution method that creates genetic diversity by recombining related DNA sequences.^33, 34^ This mimics natural recombination, creating a diverse library of variants that can be screened for improved traits such as higher enzymatic activity, increased stability, or altered substrate specificity. A key challenge limiting the large-scale use of recombinant proteins is their poor biomanufacturability. Due to toxicity issues in cellular expression, the yield of structural proteins is not high enough to be cost-effective compared to polymers, which could be addressed by high-throughput screening approaches.

In this study, we present the synthesis and high-throughput screening of a structural protein library inspired by SRT proteins. To efficiently manage the complexity of design and randomization, we use a tandem repeat library construction method based on combinatorial DNA assembly, enabling us to generate and test multiple sequence variants systematically.^35^ Our goal is to demonstrate the efficient synthesis of large libraries of structural proteins, especially those with repetitive DNA sequences. Along with our high-throughput screening platform,^36^ this can lead to the discovery of sequences that enhance biomanufacturability. The library contains over one million assembled variants from various protein fragments found in squid species. Although there are several library synthesis methods, such as random mutagenesis,^37^ directed evolution,^38^ and DNA shuffling,^39^ we chose combinatorial library synthesis to preserve native protein sequences.^26^

## RESULTS

### SRT combinatorial library

About 350 squid species are found worldwide, and we studied six of these strains (Figures 1a, b), each showing a unique SRT protein composition. These proteins have a segmented, copolymer-like structure, with fragments repeated in tandem. SRT proteins are classified as proteins with intrinsically disordered regions, where each repeat contributes to both crystalline (β-sheets) and amorphous domains (Figure 1c). In this study, we created a library of SRT proteins with four fragments that have different amorphous regions while maintaining a constant crystalline sequence. Figure 1d shows the design strategy and amino acid composition. A set of 33 amorphous sequences found in nature, with similar chain lengths (17–22 amino acids), was assembled in a combinatorial manner, resulting in 1,185,921 (33^4^) variants. SRT proteins generally tend to form β-sheets,^40^ and to enhance this native property, the crystal-forming sequence was kept constant. The predicted tertiary structures of several variants within the SRT protein library are displayed in Figure S1.

**Figure 1.**
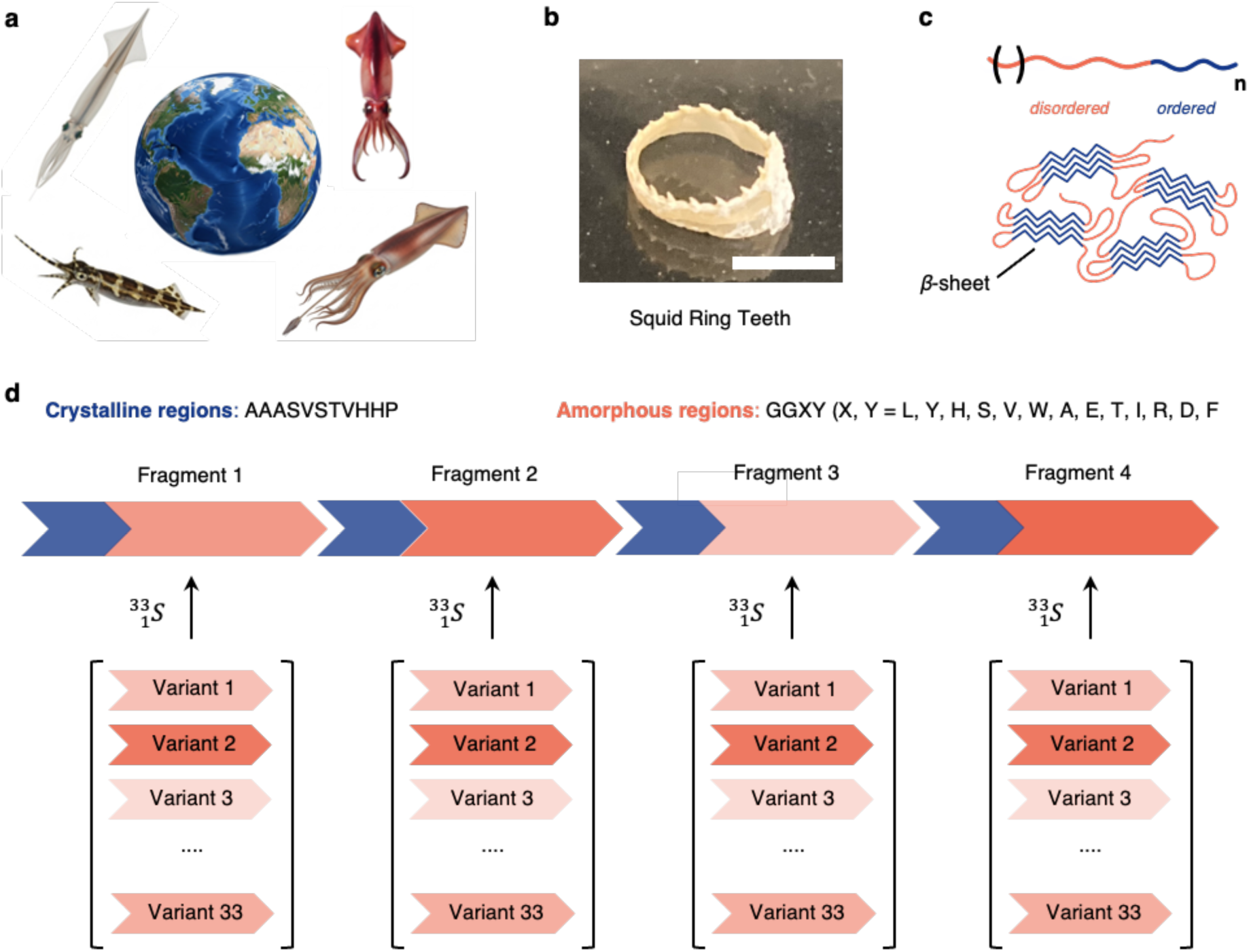
SRT protein library design. a. Sequences of SRT proteins in four squid species from around the world were studied and served as inspiration for the library design. b. A representative squid ring teeth is shown (scale bar = 5 mm). c. SRT protein repeat fragments consist of a crystal-forming region and an amorphous region, which assemble into a network of β-sheets. A protein chain is formed by ‘n’ tandem repeats. d. The design strategy of the n = 4 SRT protein library is shown. The crystalline sequence was kept constant, while a set of 33 amorphous sequences was selected from naturally occurring sequences that assemble in a combinatorial fashion.

Our library offers a comprehensive set of protein features for holistic screening and sequence-to-expression analysis. The amorphous fragments consist of different amino acids from all classifications, including polar, electrically charged side chains, hydrophobic, and special cases (Figure S2). While the overall positive hydrophobicity score of all fragments makes them hydrophobic, there is considerable variation in their degree of hydrophobicity (Figure S3). Additionally, the differences in amino acid composition cause the fragments to exhibit a range of electric charges, measured by the isoelectric point (Figure S3). It is important to note that these characteristics are amplified when four fragments randomly assemble to form a protein chain. We hypothesize that this diversity in protein building blocks affects their recombinant expression levels because of the different folding and conformational tendencies of these chains inside host cells.

### Single-cell and ensemble screening strategy

In our previous study on high-throughput protein expression optimization,^36^ we examined libraries of plasmid components, namely promoters and 5’ untranslated regions, which are known to influence single-cell protein levels. Additionally, the characteristics of these features (e.g., transcription rate, translation initiation rate, mRNA decay rate) and their subsequent effects on single-cell protein levels can now be predicted using Artificial Intelligence models. In biomanufacturability screening, it is crucial to first separate single-cell protein expression from ensemble-level expression. Since ensemble expression is affected by other factors, including growth dynamics and metabolic burden, it is more effective to establish genotype-to-phenotype correlations at the single-cell level first. Later, cellular growth characteristics can be analyzed in the context of single-cell expression to connect biomanufacturability with DNA or protein sequence. However, such predictions are not possible based solely on the protein coding sequence (CDS) because promoter strengths, ribosome binding site (RBS) strengths, and other regulatory elements were kept constant in this study. To accomplish this, we used fluorescence-activated cell sorting (FACS) to divide the SRT protein library into three logarithmically spaced bins based on mCherry fluorescence, which acts as a proxy for protein expression levels (as detailed in later sections). Cells from each bin were then sequenced to link genotype with protein expression phenotypes at the single-cell level. Our workflow is shown in Figure 2a.

**Figure 2.**
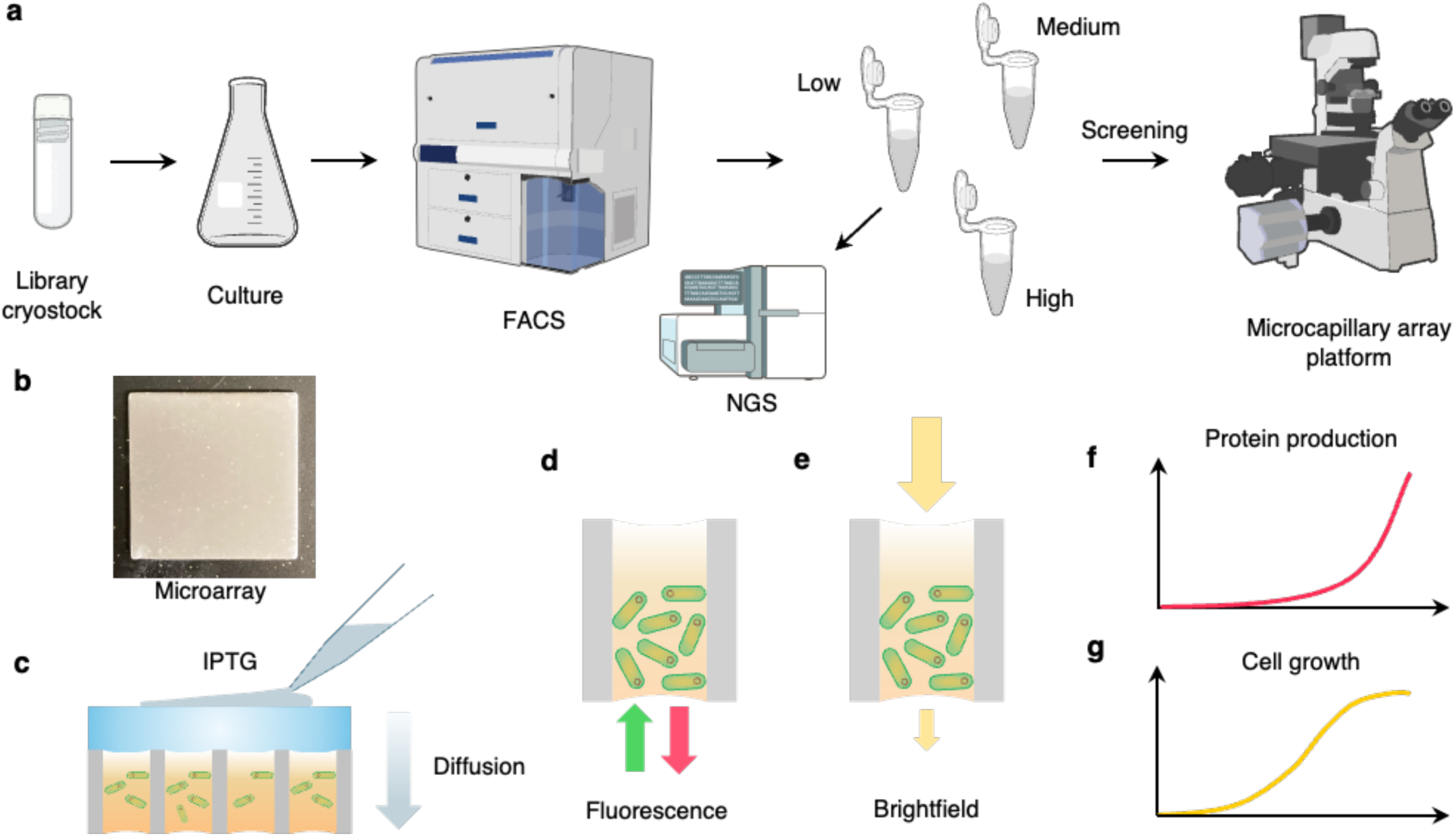
Screening Methodology for Single Cells and Ensembles. a. Single-cell protein levels are measured using cytometry, and the library is sorted into three sub-libraries—low, medium, and high—with protein levels differing by a single order of magnitude. These sub-libraries are then screened using the micro-capillary array platform to analyze growth characteristics through miniature cultures. b. A representative microcapillary array is shown (dimensions: 20 mm × 20 mm). c. For induction, stock IPTG aqueous solution is applied to the agarose gel overlaid on the array. Two optical signals—fluorescence (d) and brightfield (e)—are collected from each capillary in the array. These signals are used to estimate protein levels (f) and cellular growth (g) rates.

We then independently screened each bin’s sub-libraries using our high-throughput microcapillary array platform, shown in Figure 2b. This system enables spatially isolated clonal growth within individual capillaries (20 μm diameter) and allows for the induction of the expression system through IPTG diffusion via an agarose gel (see Figure 2c). It captures both fluorescence and brightfield signals to measure protein levels and cell density, respectively. The fluorescence intensity summed across each capillary reflects protein quantity, while the transmitted brightfield intensity is used to calculate absorbance, indicating cell density relative to a background reference. Unlike flow cytometry or plate readers, this platform supports high-throughput, time-series experiments (over 10^5^ clones per hour), facilitating the generation of production and growth rate curves over extended periods. In our workflow, the microcapillary array platform and FACS complement each other: first, by distinguishing one of the two variables critical to biomanufacturability (protein per cell or cell frequency), and second, by capturing the other variable at high resolution within the enriched search space. This approach enables a systematic and comprehensive study of how factors such as protein composition, structure, expression system design, cell viability, and environmental conditions influence biomanufacturability.

### Expression system and cloning

The plasmid design for the library is shown in **Figure 3a**. The expression system includes a T5 promoter with two lac operators, a ribosome binding site with a consistent translation initiation rate (1664 au), a protein coding sequence, a HiBiT tag, and an mCherry CDS that overlaps by four nucleotides with the stop codon of the protein CDS. SRT proteins are naturally non-fluorescent, making their direct optical screening impossible. Therefore, we use mCherry as a biomarker that is expressed in proportion to the protein CDS through ribosomal re-initiation in a bicistronic construct (**Figure 3b**).^41^ RNA polymerase first transcribes the protein CDS starting from the initial upstream open reading frame (ORF). Instead of fully disassembling at the ‘TGA’ stop codon, RNAP re-initiates translation at the downstream ORF’s ‘ATG’ start codon of mCherry. This mechanism ensures a positive correlation between the translation of both ORFs due to the short intergenic region (4 nt) and the absence of a strong transcriptional terminator.

**Figure 3.**
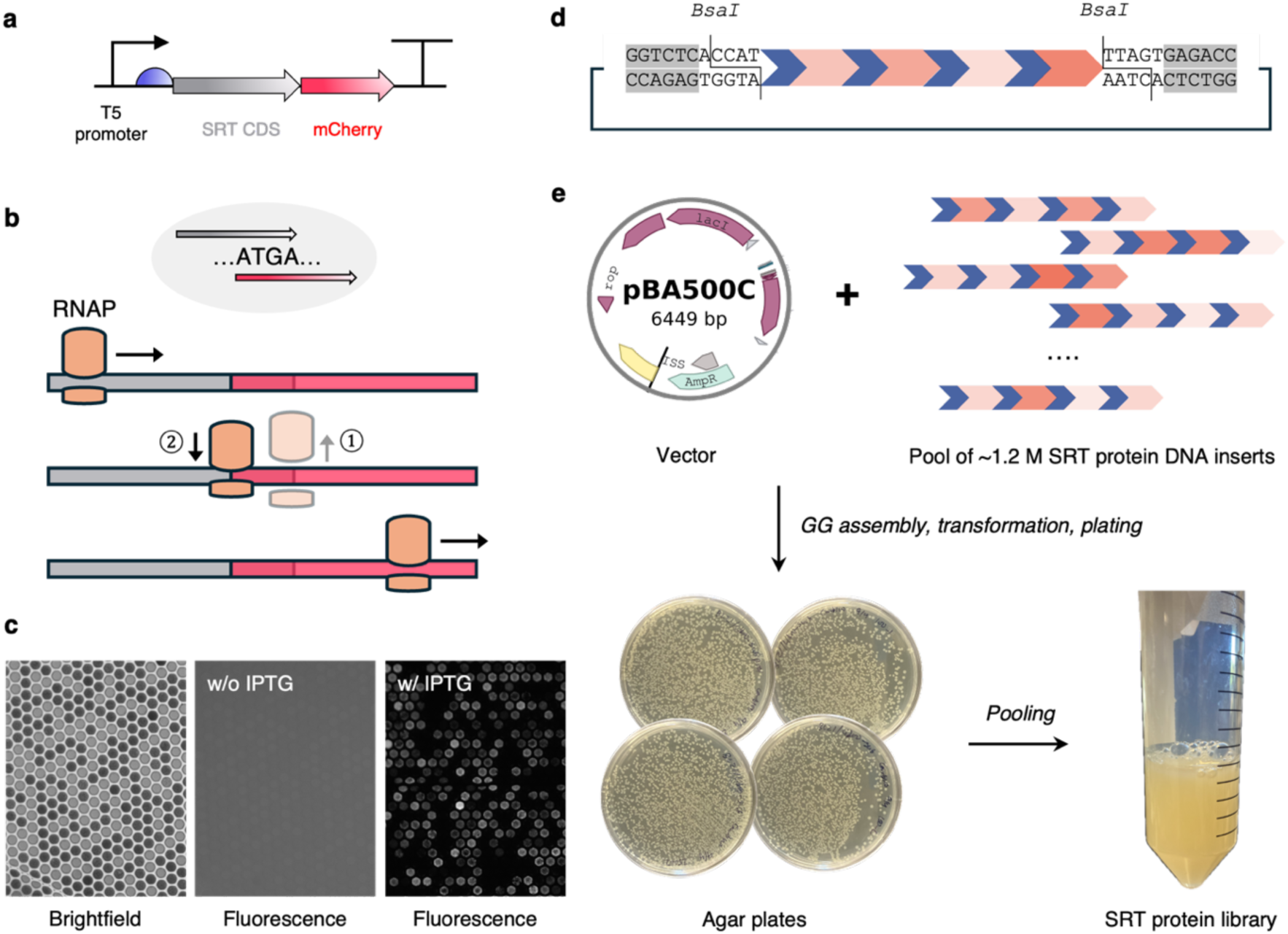
SRT protein library plasmid design and cloning. a. The SRT protein library plasmid design features key components such as the T5 promoter, ribosome binding site (shown in blue), SRT protein CDS, mCherry CDS, and T7 terminator. b. The mechanism of ribosome re-initiation is illustrated. The mCherry ORF overlaps with the SRT protein ORF by four nucleotides. c.

The T5 promoter system is recognized by native E. coli RNAP and does not require the expression of other specific polymerases. When combined with an expression control system like the lac operator, which is inducible by IPTG, T5 provides better transcriptional regulation than T7 (as shown in our previous study^36^). This helps prevent basal (‘leaky’) expression, which is especially important when expressing toxic proteins. In our vector, we included two lac operator sites near the promoter to enhance the regulation of basal expression. The effectiveness of T5’s prevention of basal expression, using the IPTG-inducible system, was confirmed with microcapillary arrays, as shown in Figure 3c. No fluorescence signal was detected in the non-induced capillaries, despite using a high exposure time (4s) for imaging. In contrast, the induced array showed significant mCherry fluorescence at about 40 ms exposure.

Images display an array of cells expressing the library plasmids. The brightfield image shows miniature cultures. No fluorescence was detected in arrays that were not induced. d. The design of the SRT protein DNA insert, sourced commercially in the pUC57 vector, is shown. The insert is flanked by BsaI restriction sites, enabling compatibility with Golden Gate assembly. e. The methodology for library cloning using the pBA500C vector and pUC57-SRT protein insert pool is illustrated.

To build the protein library, we used a streamlined cloning method with pre-assembled fragments (Figure 3d) inserted into a BsaI-flanked entry plasmid, allowing efficient Golden Gate assembly. This design significantly increased accuracy by reducing the number of assembly parts by 2.5 times. The destination plasmid pBA500C (which contains the base vector and protein insertion site) and protein-coding inserts were combined in a single reaction, simplifying construction without reducing efficiency (Figure 3e). To ensure even clone representation and reduce biases from different growth rates, we pooled isogenic colonies directly from agar plates instead of growing individual colonies in liquid culture. This method ensured that the final library maintained an unbiased composition reflecting the initial assembly.

### FACS of SRT library

Single-cell protein expression levels for the negative control and the SRT protein library are shown in Figures 4a and b. The clear separation between the two populations indicates strong mCherry expression, which is crucial for accurately identifying SRT protein-expressing clones. In addition to generally high mCherry expression, the SRT protein library covers roughly a two-order-of-magnitude range in fluorescence intensity, indicating considerable diversity in expression levels. This variation suggests that the encoded protein significantly influences expression modulation. To explore this diversity further, the library was sorted into three bins, as shown in Figure 4b, with each bin representing a specific expression range and consisting of a similar, minimal fraction (about 15%) of the total population.

**Figure 4.**
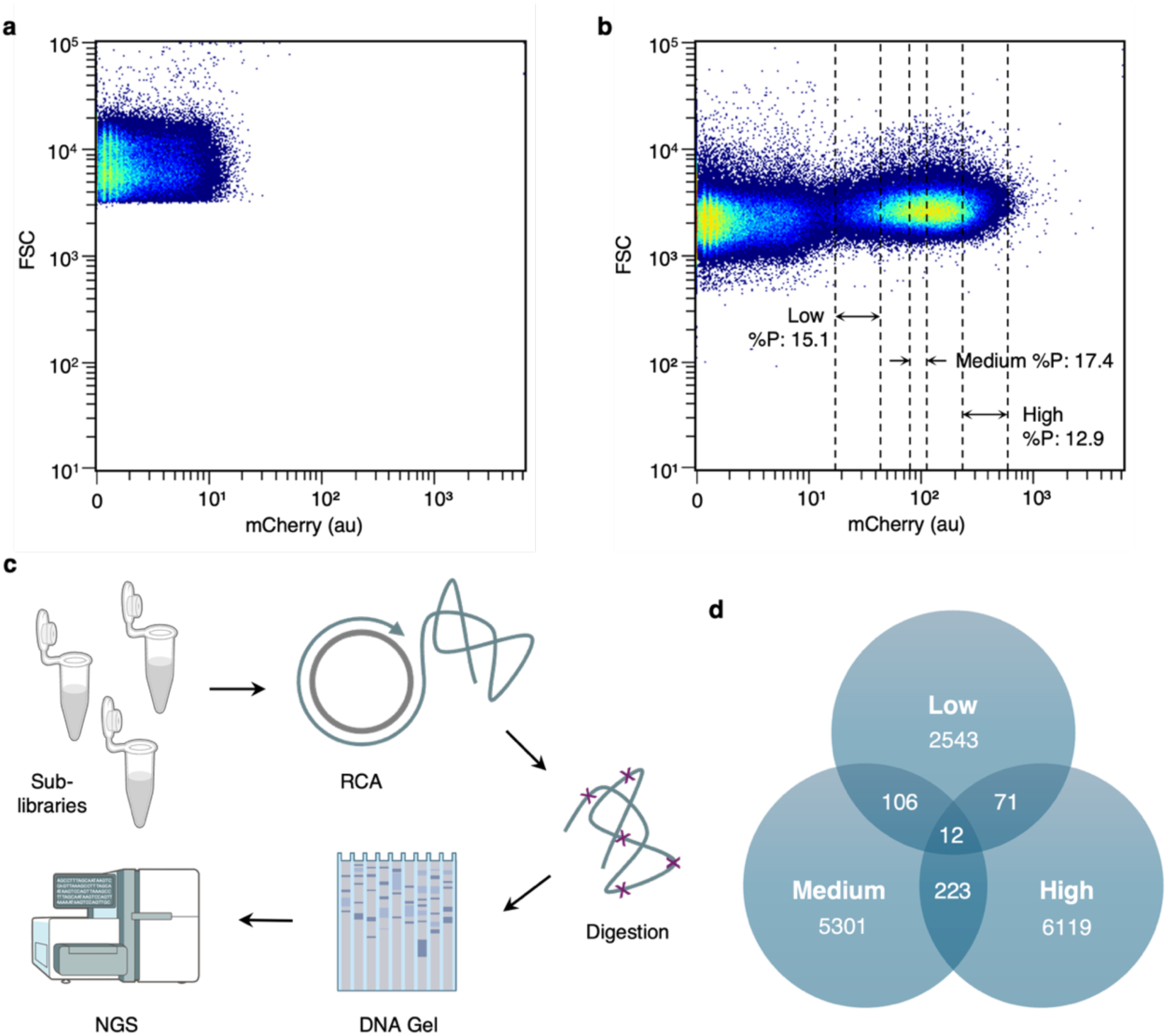
Flow cytometry distribution and FACS bins.Single-cell mCherry expression levels of the negative control (pBA500C-containing cells, a) and SRT protein library clones (b) are shown. The FACS bin gates are illustrated with their respective share of the population.

Figure 4c illustrates our cost-effective strategy for sequencing the constructs in each bin, which includes DNA template amplification through rolling circle amplification (RCA) and sequencing via Next-Generation Sequencing (NGS, Illumina). We chose RCA over polymerase chain reaction (PCR) to prevent off-target amplification in the repetitive DNA templates. NGS is ideal for short inserts like the SRT protein library, and its high accuracy (>99.9%) allows for easy and rapid alignment with the reference.^42^ Sequencing cells from each bin revealed significant diversity within each group, with minimal sequence overlap between bins (Figure 4d), further supporting the connection between sequence composition and expression behavior. The mCherry-positive density scatter plot shows that most clones express medium protein levels, with fewer clones showing low or high expression. The cell population in the sorting region resembles a unimodal symmetric distribution (Figure S4), suggesting an unbiased composition of the SRT protein library—resulting from randomized fragment assembly. This balanced distribution is ideal for establishing a strong genotype-to-phenotype relationship.

### Genotype-to-phenotype correlations

The SRT protein library includes sequence variations specifically within the amorphous regions, resulting from differences in amino acid composition and fragment length. To explore potential relationships between genotype and phenotype, we analyzed correlations between various protein characteristics and single-cell expression levels. Specifically, we compared hydrophobicity (Wimley–White scale), isoelectric point, molecular weight difference, and nitrogen content of sequences in the low, medium, and high bins. No statistically significant correlations were found for any of these properties. Importantly, features of the expression-regulating plasmid, such as the promoter and ribosome binding site, were kept constant across all variants in the library, ensuring that differences in expression were solely due to the encoded protein sequences.

Since single-cell expression seems unaffected by the intrinsic physicochemical properties of the proteins encoded, we examined sequence features that could influence folding efficiency— specifically, intra-sequence self-similarity. We conducted pairwise comparisons of 17–20 amino acid fragments (see Figure 5a for workflow), generating multiple pairs for each bin to ensure reliable statistical analysis. The distribution of fragment-pair identities was unimodal and symmetric, indicating that the combinatorial library is unbiased (Figure 5b). FACS analysis revealed clear enrichment patterns: low-expression bins were enriched with fragment pairs sharing 10–30% identity, with most sequences containing one or fewer such pairs (Figure 5c). In contrast, high-expression bins favored higher self-similarity, with 40–60% identity pairs being most common, and most sequences containing three or more pairs with over 50% identity. The medium bin’s similarity levels were intermediate. This indicates that higher intra-sequence homology is linked to, and may promote, increased protein expression in our system.

**Figure 5.**
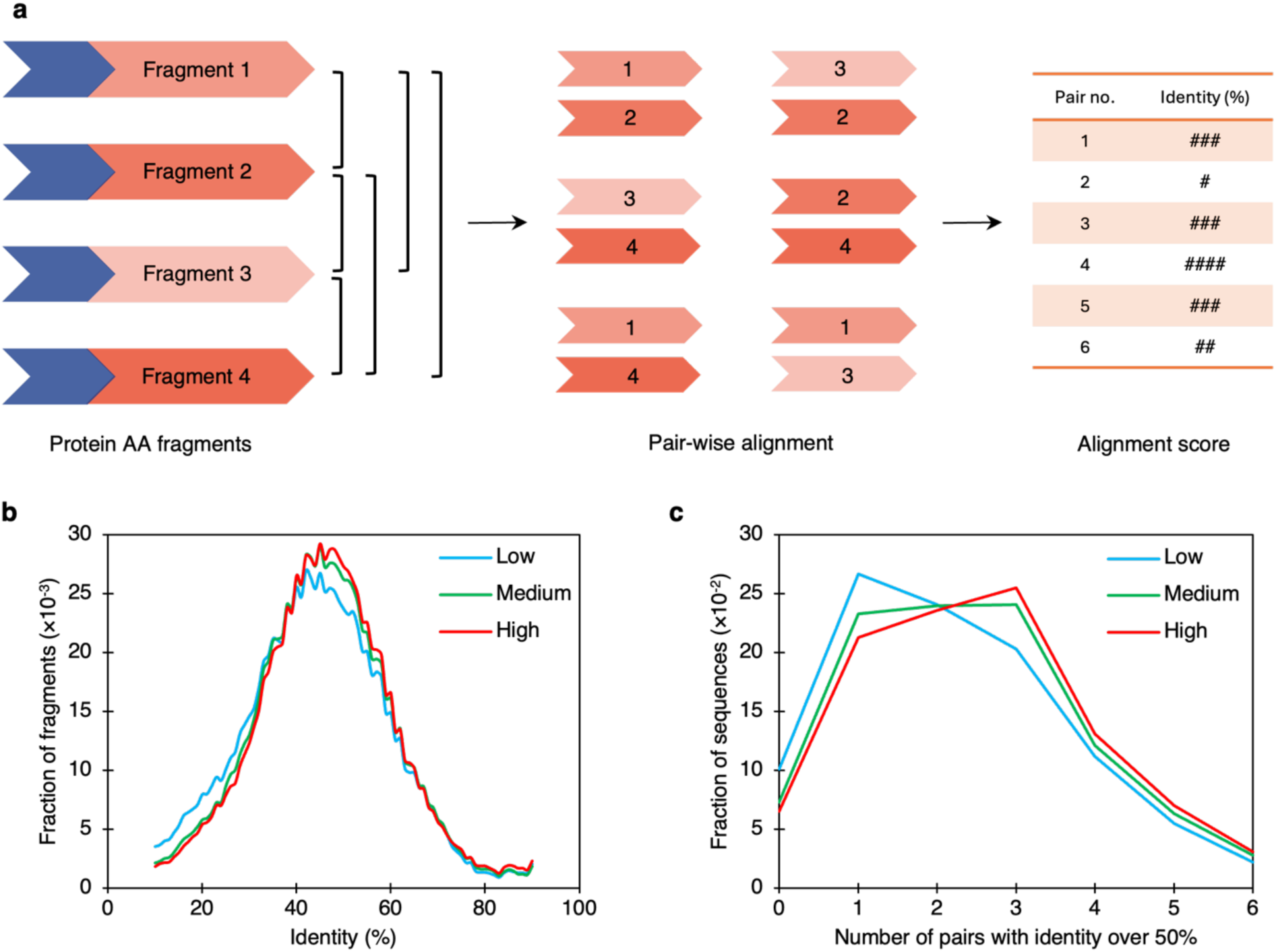
Analysis of sequence self-similarity in SRT library. a. The strategy for checking sequence self-similarity among fragments in protein chains is illustrated. For each protein chain, pairwise alignment of six pairs of fragments is performed, and percent identity is recorded. b. Distribution of percent identity in fragment pairs within each bin is plotted. c. Distribution of sequence population versus the number of pairs with over 50% identity is plotted.

### High-throughput screening using microcapillary arrays

We expanded the orthogonal screening of the SRT protein library by utilizing our high-throughput platform, which produces protein and growth data that traditional methods cannot achieve. We analyzed the bins over a 35- hour period, measuring mCherry fluorescence levels (Figures 6a, b, c) and cell density (Figures 6d, e, f) in more than 15,000 capillaries in a series of time points. Not all capillaries exhibit the same mCherry fluorescence, highlighting significant heterogeneity in protein expression. The fluorescence intensity distribution is broad, a common trait of stochastic gene expression. This variability is particularly evident in structural proteins, which are generally more challenging to express compared to soluble fluorescent proteins (as also observed previously with cement and reflectin versus mRFP1^36^).

**Figure 6.**
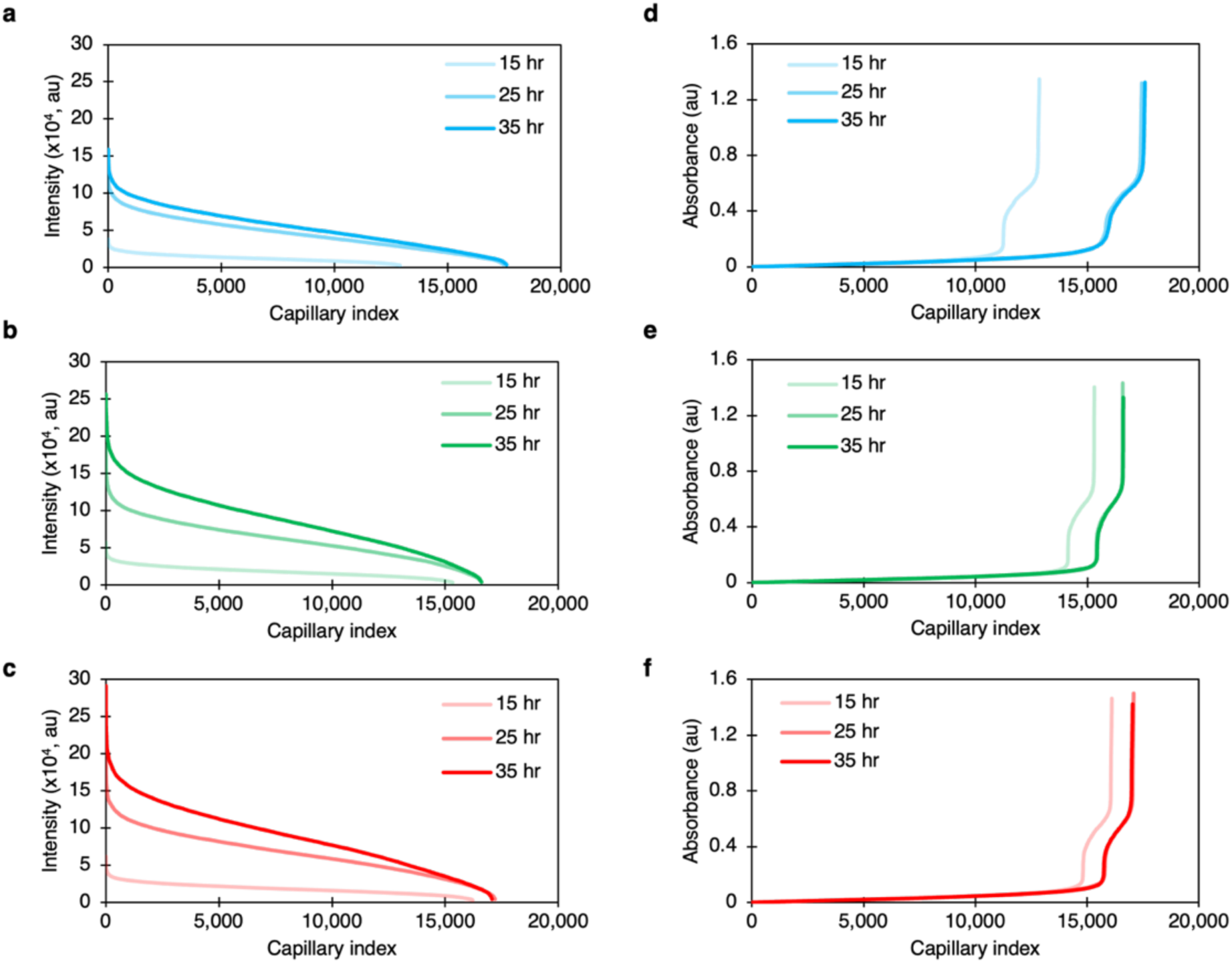
Protein production and cellular growth rates. Time series plots of protein production (a, b, c) and cell growth (d, e, f) for the low (a, d), medium (b, e) and high (c, f) bins at different time points after induction.

Interestingly, the distribution is asymmetric — it appears skewed, with a sharp increase in the number of capillaries at high fluorescence intensities and a gradual decline toward lower intensities. This "pinning" at the lower end likely reflects a combination of expression inefficiencies and biological factors such as cell viability. Indeed, in each expression bin, a subset of clones produced minimal levels of protein, which could be due to compromised cell health or other cellular burdens. Moreover, the rate at which intensity values increase in each bin provides additional insights (Figure S5). The rise in intensity from t = 15 hr to 25 hr is greatest in the high bins, followed by medium and low. Additionally, the low bin showed minimal additional expression during the final 10-hour incubation period. The medium and high-expression bins show a broader, more gradual increase in fluorescence values. This indicates a broader range of expression levels and potentially more resilient or variable cellular states within those populations. Fluorescence images served as templates to identify and estimate the number of bright capillaries, which were then matched with an equal number of the darkest capillaries in the corresponding brightfield images to measure protein absorbance. Consistent with this, the number of capillaries exceeding the low bin’s threshold in both the medium and high bins was significantly higher (as shown in Figure 6a and 6d). This trend highlights the underlying diversity in expression potential across the library and supports the existence of a dynamic, non-uniform distribution of expression phenotypes.

Representative fluorescence and brightfield images are shown in Figures 7a and **7b**, respectively. The resulting protein production and cell growth profiles at the final time point are summarized in Figures 7c and **7d**, respectively. A clear upward trend in protein titer is observed across the expression bins—from low to medium to high—with each bin exhibiting a higher titer than the previous one. Notably, the medium-expression bin displayed approximately 54% higher protein titers compared to the low-expression bin. In contrast, the increase from the medium to high bin was modest, with the high bin showing only about a 5% improvement over the medium bin (Figure 7e).

**Figure 7.**
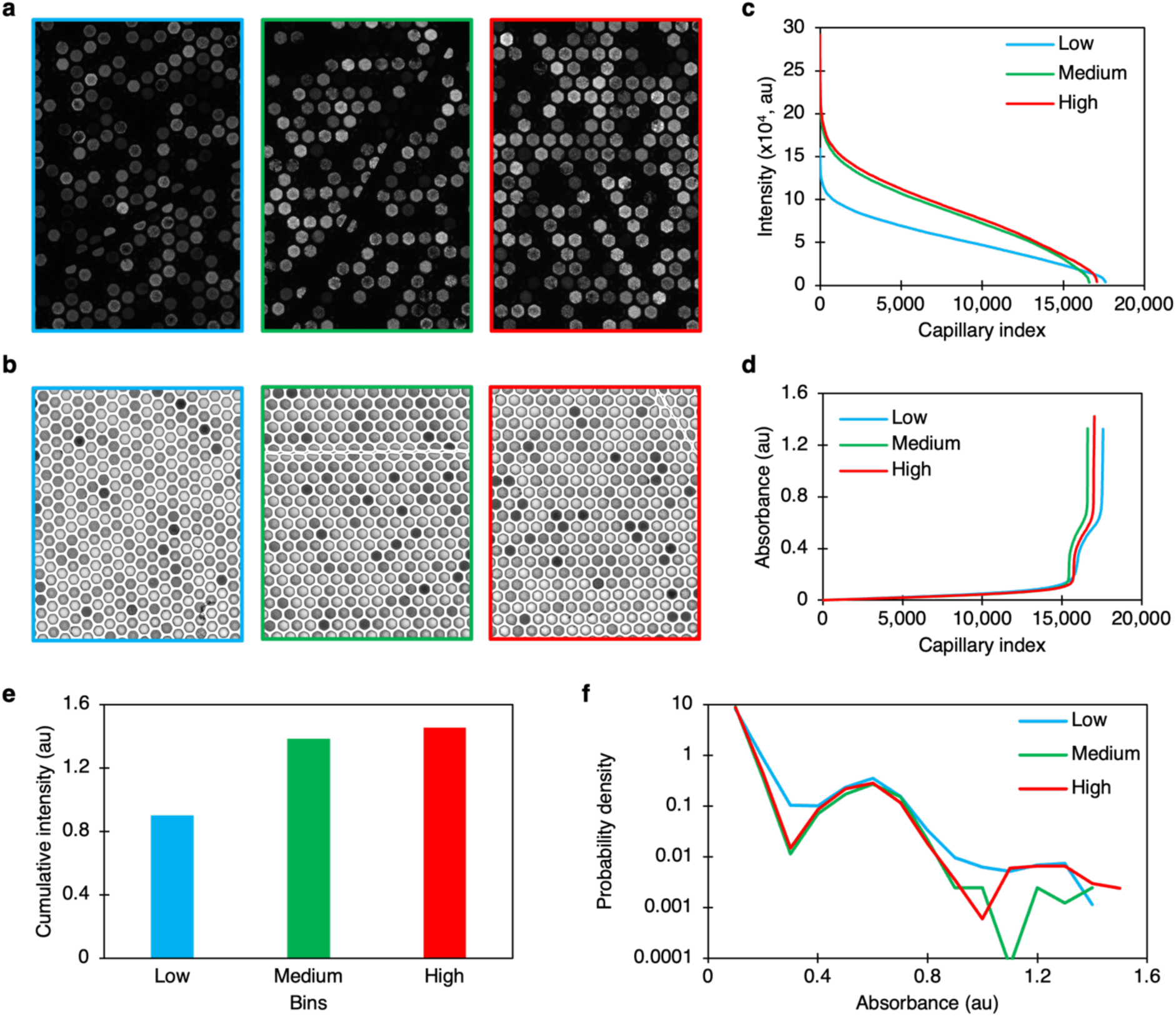
Comparative analysis of protein production and cell growth. Fluorescence (a) and brightfield (b) images showing capillaries for the low (blue), medium (green), and high (red) bins at the t = 35-hour time point. Protein production (c) and cell growth (d) curves at t = 35 hours for the bins are compared. e. Cumulative fluorescence intensities of the bins at t = 35 hours are shown. f. Distribution of capillary frequency versus absorbance at t = 35 hours is depicted.

The platform’s main insight is that growth rates are similar across all bins. Although it was expected that clones in the high bin might grow slower due to metabolic burden from accumulating more protein per cell, and that low and medium clones might grow better, all three bins showed nearly identical absorbance (or cell density) distributions at t=35 hr (Figure 7f). This distribution also matches that at t=15 hr for each bin, indicating that the cell population in the capillaries stabilized early on—probably during the pre-induction phase at 37°C—and then remained stable during the later incubation at 25°C after induction. This stability is likely due to the T5 expression system’s ability to suppress basal expression and the lower temperatures used after induction , which help cells grow largely independent of the expressed protein.

These results directly correlate with the single-cell protein levels; however, the same degree of variation observed in the single-cell protein levels was not seen at the ensemble level of protein expression. Each clone is expected to show a range of single-cell protein levels, and selecting cells with the average protein level is not guaranteed in FACS, especially for the high bin, making a 10- fold variation in protein titers unlikely. Additionally, unviable cells are present in each capillary, as confirmed by cytometry of individual cultures from several clones in the library (Figure S6) and the dense left lobe in Figure 4b. Therefore, it is likely that only a portion of the clonal population expresses mCherry and SRT proteins (as previously noted with cement and reflectin ), and this fraction varies across the bins. Our results suggest that clones in the medium bin perform similarly to those in the high bin, possibly because they compensate for lower single-cell protein levels with a larger number of cells expressing the proteins (illustrated in Figure S4). The mCherry-positive cell populations appear to be centered around the medium bin; the sharp increase in cell frequency toward the high bin supports this as well (Figure S4). However, further studies are needed to definitively determine the reasons, which is beyond the scope of this study.

## DISCUSSION

SRT proteins are recognized for their multifunctional properties, making the optimization of their biomanufacturability essential for practical applications. While previous studies have examined how protein sequence affects biomanufacturability in various systems, our research expands this investigation to include SRT proteins and broadens its scope by utilizing an extensive combinatorial library. This library was designed to cover a wide sequence space, including variations in features such as self-similarity, hydrophobicity, charge, and nitrogen content. Significantly, sequences responsible for crystalline region formation were kept constant to ensure structural consistency across variants. Our high-throughput screening used a microcapillary array platform capable of capturing the entire distribution of production levels within each bin, stratified by single-cell expression intensity. This enabled a detailed mapping of genotype to phenotype across thousands of clones. The platform, based on T5 promoter-driven expression, effectively minimized basal expression, resulting in uniform growth profiles across bins. A strong correlation was found between single-cell protein expression and final protein titer levels. However, within the high-expression bin, a wide range of titer values was observed, emphasizing the influence of stochastic expression and cell viability constraints even among high-expressing clones. Notably, the platform features a precision extraction mechanism that allows for iterative enrichment of the high bin. This makes it possible to progressively isolate top-performing clones through multiple rounds of screening and refinement, providing a powerful tool for engineering biomanufacturable variants at scale.

While most physicochemical properties of the protein sequences—such as hydrophobicity, charge, or nitrogen content—were not found to strongly correlate with protein titer levels, sequence self-similarity emerged as a key factor influencing expression. SRT proteins consist of tandem repeats that form β-sheet structures. This assembly behavior is reflected in their inherent disorder profiles. For example, the engineered variant TR-n4, made up of four tandem repeat units, is known to form β-sheets and displays a consistent, periodic disorder profile with four distinct peaks (Figure S8). These peaks indicate flexible regions in the sequence that undergo structural rearrangement during folding. To explore the connection between self-similarity and β-sheet assembly, we analyzed the disorder profiles of the most self-similar protein sequences in our library (cumulative self-similarity score: 260 – 600) and compared them to the least self-similar sequences (score: 266 – 273). The results, shown in Figure S9, reveal that highly self-similar clones display periodic disorder patterns with four evenly spaced peaks, closely resembling the TR-n4 profile. In contrast, the disorder profiles of low-similarity sequences are irregular and lack clear periodicity. These findings suggest that sequence self-similarity encourages regular folding into β-sheet structures, thereby increasing expression levels. However, further investigation is needed to understand how individual chain disorder influences protein-protein interactions, which could be directly related to aggregation tendencies and protein toxicity in E. coli. Despite maintaining consistent crystal-forming sequences, the amorphous regions appear to impact self-assembly propensity. This creates a mechanistic connection between primary sequence design and the biomanufacturability of SRT proteins.

In summary, we have demonstrated an orthogonal screening method that combines FACS and fluorescent microcapillary arrays to establish genotype-to-single-cell-to-ensemble correlations. We observed that brighter clones tend to contain self-similar sequences (i.e., perfect repeats of fragments), whereas dimmer clones in the library have less similarity (i.e., imperfect repeats). The method employs two high-throughput techniques in a complementary way, covering screening from single cells to the ensemble level, while enabling enrichment of desired clones. Our workflow separates single-cell expression and growth traits for a systematic study of genetic design on biomanufacturability, and is especially suited for genetic systems that do not respond well to computational predictions, such as those involving structural proteins.

## DATA AVAILABILITY

The authors declare that data supporting the findings of this study are available within the paper and its supplementary information files.

## AUTHOR CONTRIBUTIONS

M.C.D. conceived the project. T.M.B. and K.S. developed the platform. B.D.A. designed the plasmids. K.S. performed the cloning, designed the experiments, and conducted screening. K.S. wrote the manuscript together with M.C.D. All authors participated in manuscript revisions, discussions, and data interpretation.

## ACKNOWLEDGMENTS

We thank Dr. Howard Salis and Harry Adamson for their scientific discussions. Additionally, we appreciate Dr. William Humphries for guiding us in setting up the optical system.

## ACKNOWLEDGEMENT OF SPONSORSHIP STATEMENT

This effort was sponsored in whole or in part by the Central Intelligence Agency (CIA), through CIA Federal Labs. The U.S. Government is authorized to reproduce and distribute reprints for Governmental purposes notwithstanding any copyright notation thereon.

## DISCLAIMER

The views and conclusions contained herein are those of the authors and should not be interpreted as necessarily representing the official policies or endorsements, either expressed or implied, of the Central Intelligence Agency.

## METHODS

### Plasmid library design

The amino acid sequences of the protein are based on native Squid Ring Teeth proteins.^21^ We designed a segmented copolymer protein with four fragments. Each fragment has a crystalline, beta-sheet forming region and a flexible, GGXY amorphous region. The amino acid sequence of the crystalline region across all fragments remained constant at ‘AAASVSTVHHP’. We selected 33 amorphous amino acid fragments from six different squid species, based on our previous analysis. The selection was made according to the similarity in length to the original recombinant n4 sequence, as described in our earlier publication.^21^

### Vector cloning and library synthesis

Plasmids pBA407 and pBA500B were used to clone the pBA500C vector. Plasmid pBA500B containing the protein insertion site, GS linker, HiBiT tag and mCherry CDS was obtained commercially (Genscript). pBA407 was digested at BsaI and NcoI restriction sites and pBA500B was digested at KpnI and BamHI restriction sites using the standard protocols (New England Biolabs, NEB). The digested products were then assembled to form pBA500C through standard HiFi DNA assembly (NEB) and transformed into NEB 10-beta at 37°C. Isogenic colonies were propagated in Luria Broth (LB) at 37°C and 250 RPM, and the cultures were miniprepped to obtain the purified pBA500C plasmid. pBA500C sequence was confirmed through long-read whole-plasmid sequencing (Oxford Nanopore, Quintara Biosciences). pBA500C plasmid contains the SRT protein CDS insertion site compatible with Golden Gate assembly. Plasmids containing the protein CDS (flanked with BsaI restriction sites) of all variants were ordered as a pool in a commercial vector pUC57 (Genscript). Golden Gate assembly of pBA500C and protein-insert plasmid pool were carried out using standard procedures (NEB, BsaI-HF v2). The assembled product was transformed into NEB 10-beta chemically competent cells. Multiple agar plates were prepared using different loading volumes of the recovery (2.5 μl, 25 μl, 100 μl, and 250 μl). Three plates with the greatest density of isogenic colonies were selected for library pooling, and all colonies were collected using LB. A portion of the recovered colony suspension was miniprepped. The miniprepped plasmids were then transformed into *E. Coli* BL21-DE3 (NEB) chemically competent cells. All colonies from four agar plates with 100 μl and 250 μl loading volume of the recovery were collected using LB and pooled. This pool was stored in glycerol cryostocks and was used for further screening and tests. All nucleotide sequences used in this study are provided in the Supplementary Information.

### Instrumentation and sample preparation

The platform was described in detail earlier.^36^ Briefly, an inverted fluorescence microscope (Olympus IX73) was equipped with a monochrome camera (DP23M, Olympus), a motorized XY stage (Marzhauser Wetzlar Tango 3), a motorized focus drive (Marzhauser Wetzlar), and a 10x objective lens (Olympus). The platform was operated using the cellSens (Olympus) software suite. Microcapillary arrays with 20 µm capillary diameter (INCOM, Inc.) were used throughout all experiments, each containing approximately 8×10^5 capillaries. The arrays were sterilized prior to use and treated with a plasma wand to increase hydrophilicity. The cryostock of each bin was incubated in carbenicillin-supplemented Luria broth (LB) at 37°C until reaching an optical density of about 0.1. Cells in LB were loaded onto an array, with cell concentration adjusted so that each capillary typically contained a single cell. After loading, the array was overlaid with a 2% (w/v) agarose gel layer (1–2 mm thick) and incubated at 37°C in a sealed petri dish lined with moist wipes for 5 hours. The cells were then induced with IPTG by adding the stock solution to achieve a 400 μM concentration in the capillaries. Following induction, the arrays were incubated at 25°C and screened at 5, 15, 25, and 35 hours. After screening, the arrays were sterilized in 70% ethanol for several hours. They were then thoroughly cleaned with a stream of DI water, with intermediate ultrasonication for 2 minutes.

### FACS

The library cryostock was inoculated in carbenicillin-supplemented LB and incubated at 37°C and 250 RPM for 5 hours. The culture was then induced with IPTG at a final concentration of 400 μM and incubated overnight at 25°C. The culture was diluted 500× in phosphate-buffered saline (PBS, 1×) and used for FACS (ThermoScientific, Bigfoot Cell Sorter). The parameters for the gates (low, medium, and high) and the respective populations are described in the main text and figures. The sorted cells (10,000 in each bin) were collected in carbenicillin-supplemented LB and incubated overnight. Finally, glycerol stocks of each bin were prepared and used for further analyses.

### Amplification and sequencing

To obtain protein CDS inserts for sequencing, the BL21-DE3 sub-libraries were propagated in LB (37°C, 250 RPM) and stored in aliquots. Rolling circle amplification (RCA) was performed with these aliquots using standard protocols (phi29-XT RCA kit, NEB) for 24 hours. Initial verification of amplification used random primers; after confirmation, sequence-specific primers (5’-CGTGAAACATC*C*T*G-3’, 5’- GTCCTGAAGAG*A*G*G-3’) minimized background. The RCA products were diluted twofold and digested with MscI and NcoI. The digested products were run on DNA purification gels (1% w/v agarose in TAE buffer) for about an hour. Desired bands were excised and purified using a DNA gel purification kit (NEB). Plasmid concentrations were quantified with Qubit and sent for sequencing via next-generation sequencing (NGS, Illumina; Quintara Biosciences). A custom MATLAB script was used to analyze the merged raw reads.

### Microcapillary array platform screenin

Brightfield and fluorescence images were captured at t = 5 hr, 15 hr, 25 hr, and 35 hr at multiple locations across each bin array. These images were taken with consistent exposure times (450 ms for brightfield and 100 ms for fluorescence). Fluorescence images were processed using a custom MATLAB script. The contrast was first normalized, and the images were binarized to highlight white capillaries indicating fluorescence. Capillary regions were identified and used as masks to measure the total pixel intensity within each capillary in the original images. No fluorescence was detected at t = 5 hr. The threshold level used for the low bin images at t = 15 hr was applied to all other images, including medium and high bins. Similarly, brightfield images were binarized using an optimal threshold, capillaries identified, and their intensities recorded. The intensities from each image were sorted in ascending order, and the lowest set of capillary intensities—matching the number of capillaries identified in the corresponding fluorescence images—were used to generate growth curves. Remaining capillaries were classified as background. Absorbance levels were also calculated.

### Genotype-to-phenotype correlations

The workflow for calculating the self-similarity of SRT sequences is as follows: (i) For each complete amino acid sequence, extract the four variable (amorphous) fragment sequences; (ii) Calculate the six different pairwise sequence alignments among the four fragments; (iii) Compute the percent sequence identity for each pair of aligned fragments (resulting in six percentage values per sequence); (iv) Compare the low, medium, and high fluorescence bins based on the distributions of these sequence identity values.Figure 5b shows all pairwise alignment scores based on the workflow described above. The number of scores in each bin is six times the number of sequences in that bin. As you move from the Low bin to the High bin, lower-identity scores (10-30%) become less common, while higher-identity scores (40-60%) become more frequent. Note that the histogram is smoothed by averaging over windows of width 20 around each point, which explains why identities below 10% and above 90% are not displayed. Figure 5b analyzed the identity scores as a group, without considering how higher or lower pairwise identity scores are distributed within individual sequences, which we focused on in Figure 5c. The steps for these calculations are: i) Choose a cutoff value (in this case 50% sequence identity), and determine how many of the six pairwise alignments per sequence exceed this cutoff; ii) Moving from the Low bin to the High bin: fewer sequences have only one alignment (or fewer) that meets the cutoff, while more sequences have three or more alignments that meet the cutoff. We note that similar trends are observed with other cutoff values (35%, 40%, 45%).

