## supplemental figures and tables for "Biomanufacturability of Squid Ring Teeth Protein Library via Orthogonal High-Throughput Screening"

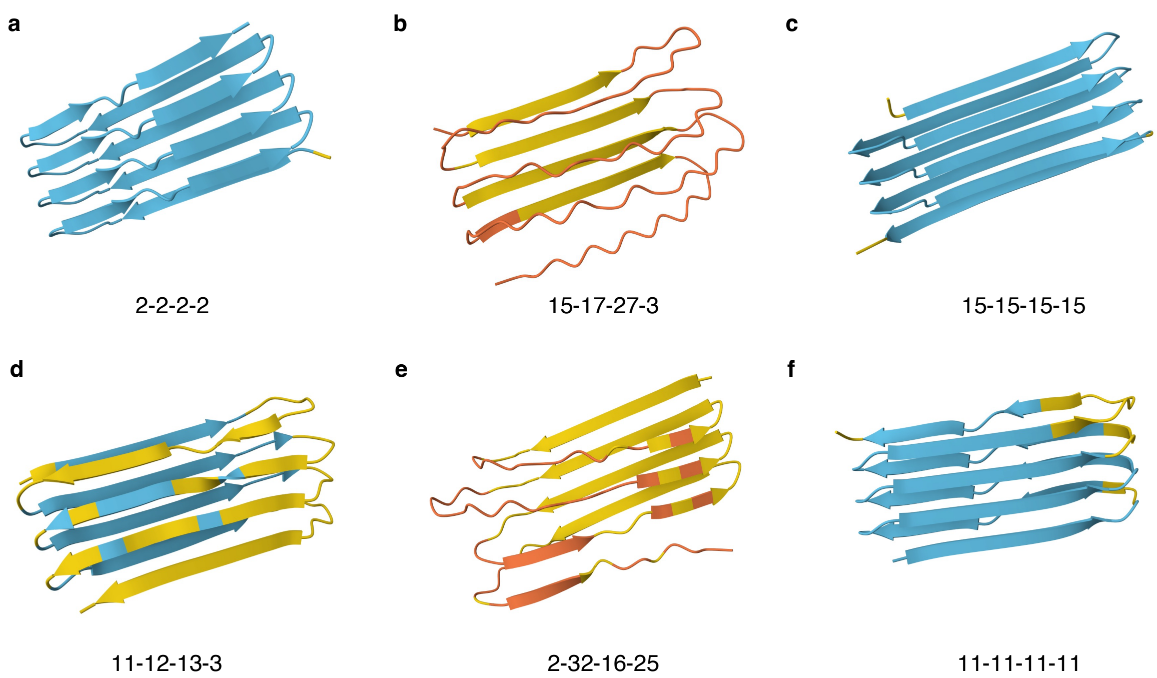


**Figure S1. SRT protein structure predictions.** AlphaFold 3 predictions of several homogenous (**a, c, f**) and heterogenous (**b, d, e**) SRT protein variants in the library. Homogenous (or self-similar) variants show high-confidence, perfect β-sheet conformations. Color-code represent standard AlphFold 3 pIDDT levels.

**a**

| | |
|---|---|
| 1 | GLGLGYGYGLGHGLGG |
| 2 | GYGLGLGLGGAGYGYG |
| 3 | GYGLGLGYGLGLGAGG |
| 4 | GYGYGGLLGGYGLHYG |
| 5 | VGYGGFGLAGYGYGYG |
| 6 | YGLAGYGGLYGGLLHG |
| 7 | YGYGGLYGGLYGGLGG |
| 8 | YIGRSVSTVSHGSHYG |
| 9 | ALGGYGGYGLGGIVGGG |
| 10 | GLGLGYGLGLGYGYGVG |
| 11 | GYGYGGLLGGLGHAVGG |
| 12 | HADYGVSGLGGYVSSYG |
| 13 | YGYGGLYGGLYGGLGYG |
| 14 | ALGEYGGYGLGGIVGGHG |
| 15 | HLGLGLGYGYGLGHGLGG |
| 16 | VGYAGYGLGLYGAGYHYG |
| 17 | YAYGGLYGGYGLGAYGYG |
| 18 | YGGIGLGGLYGGYGAHFG |
| 19 | YGYGLAGYGGLYGGLLHG |
| 20 | YSYGGLVGGYGGLYHHAG |
| 21 | AGLGYGLGGVYGGYGLHAG |
| 22 | GYGGYGLGFGGLYGGFGYG |
| 23 | GYGGYGLGLGGLYGGLHYG |
| 24 | GGYGSLLGGHGGLYGGLGLG |
| 25 | HSVSYGGWGFGHGGLYGLHG |
| 26 | LGYGGLLGGYGGLHHGVYGG |
| 27 | VGGYGGFGLGGYGGYGLGGG |
| 28 | VGYGGFGLGFGGLYGGLHYG |
| 29 | VLSGGLGLSGLSGGYGTYRG |
| 30 | YGDVYGGLYGGLYGGLLGAG |
| 31 | GFGGYGLGGYGLGGYGLGGYG |
| 32 | GYGGLYGHYGGYGLGGAYGHG |
| 33 | GYGGWGYGLGGWGHGLGGLGG |

**b**

| Category | Amino acids |
|---|---|
| Polar | S, T |
| Hydrophobic | L, Y, V, W, A, I, F |
| Electrically charged | H, E, R, D |
| Special | G |

**c**

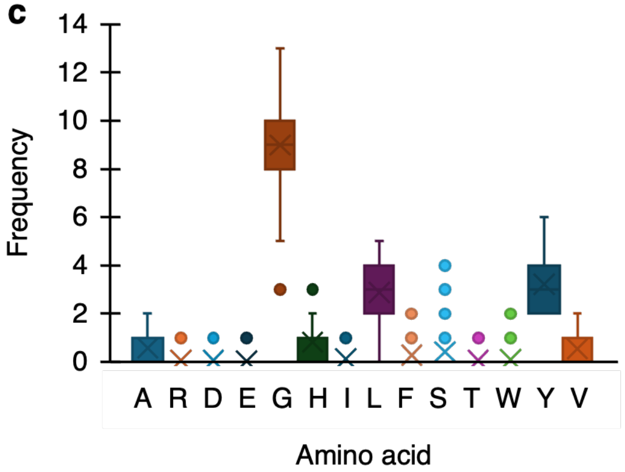


**Figure S2. SRT library fragments and amino acids a.** List of amorphous region fragments designed to synthesize the SRT library. **b.** List of amino acids utilized in the fragment pool and their classification. **c.** Frequency chart of amino acids in the fragment pool.

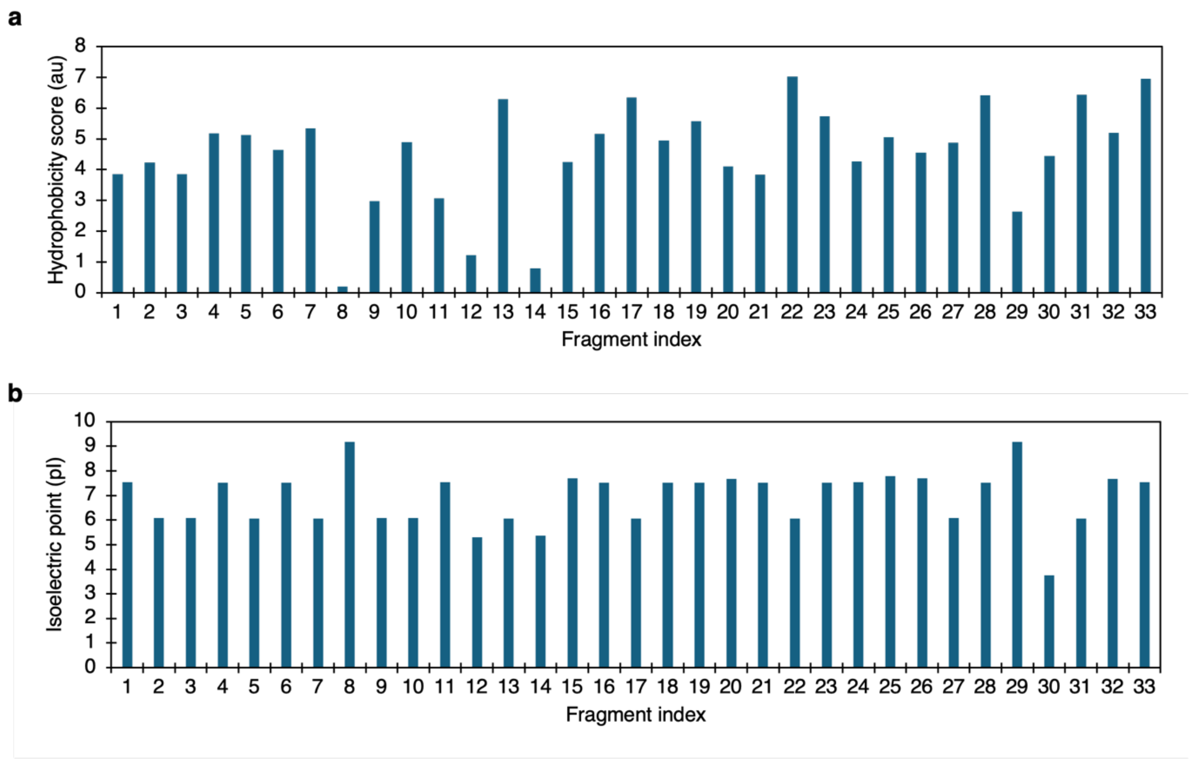


**Figure S3. Hydrophobicity and isoelectric point.** Plots depicting hydrophobicity scores (**a**) according to the Wimley-White scale, and isoelectric points (**b**) of the designed fragments.

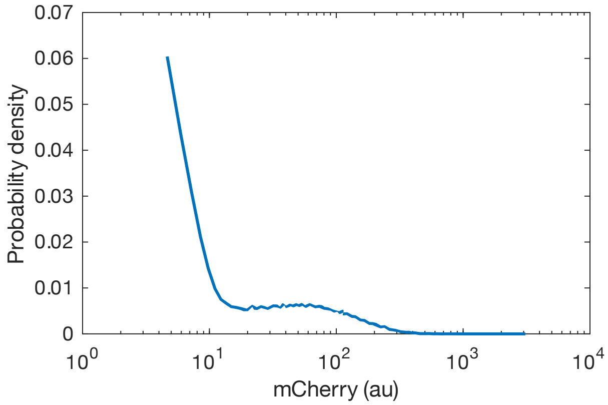


**Figure S4. Flow-cytometry frequency chart.** Distribution of single cell clones in the SRT library with respect to mCherry fluorescence.

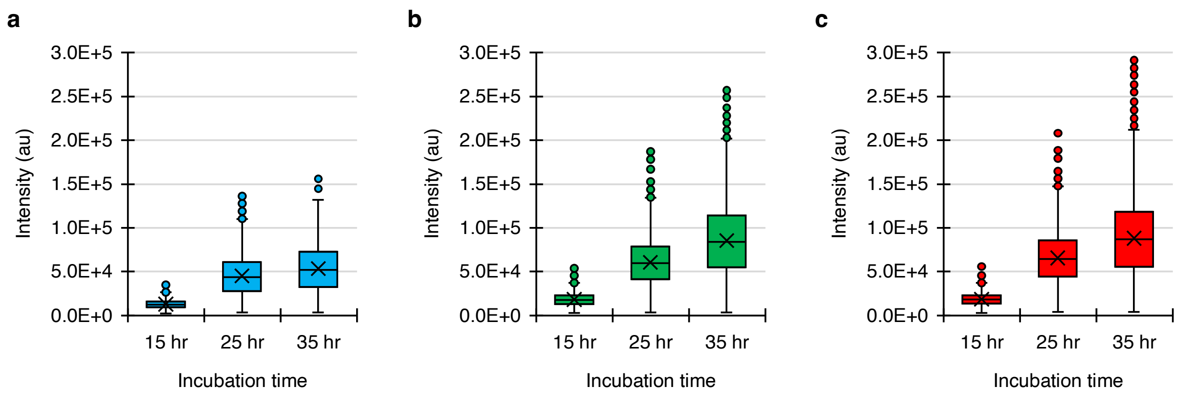


**Figure S5. Time-series protein production charts.** Distribution of fluorescence intensities of capillaries containing clones from the low (**a**), medium (**b**), and high (**c**) bins.

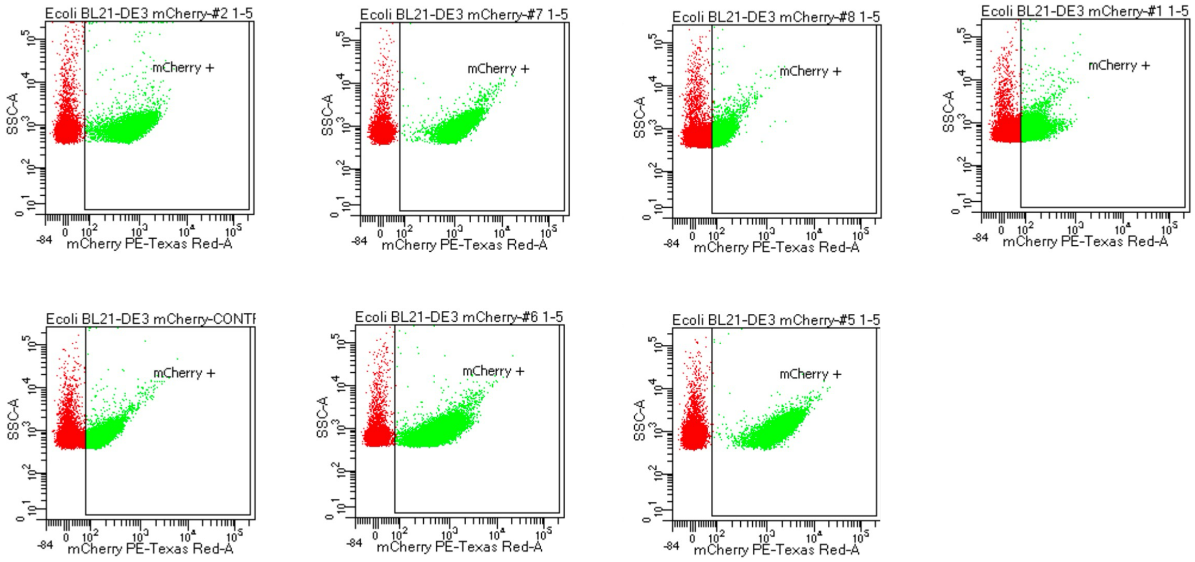


**Figure S6. Flow cytometry charts of individual clones.** Flow cytometry plots illustrating single cell mCherry fluorescence levels of several clones from the SRT library.

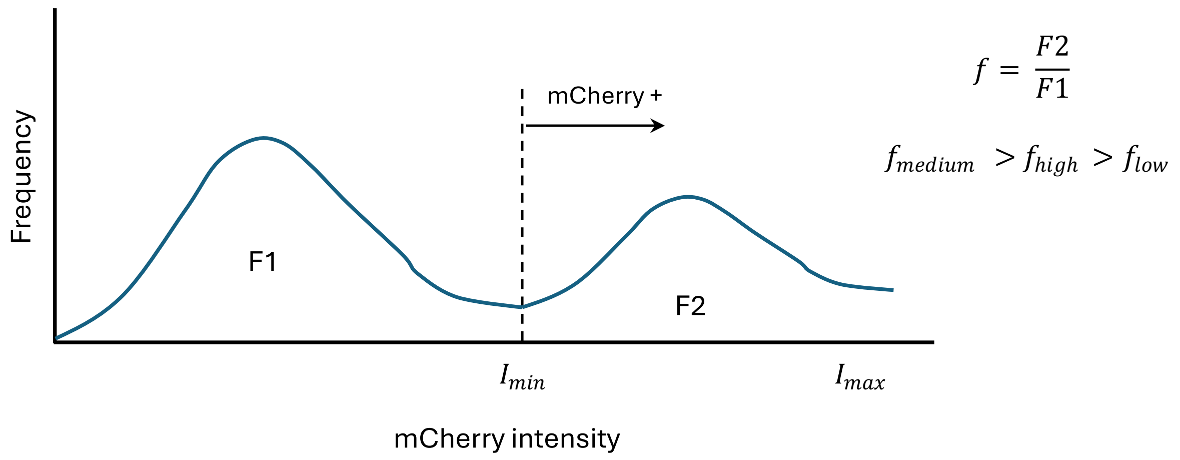


**Figure S7. Distribution of single-cell fluorescence.** The schematic describes the postulated distribution of cells in the library, where only a fraction of cells per variant express mCherry. This fraction is greater for the medium bin, enabling it to offset the loss in protein titer due to lower single cell expression.

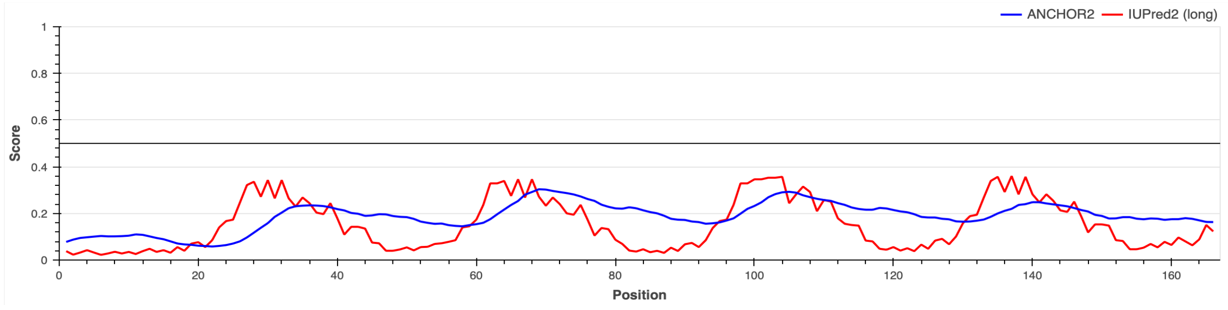


**Figure S8. Disorder measurements.** Periodic disorder profile with four distinct peaks measured by protein disorder levels (https://iupred2a.elte.hu)

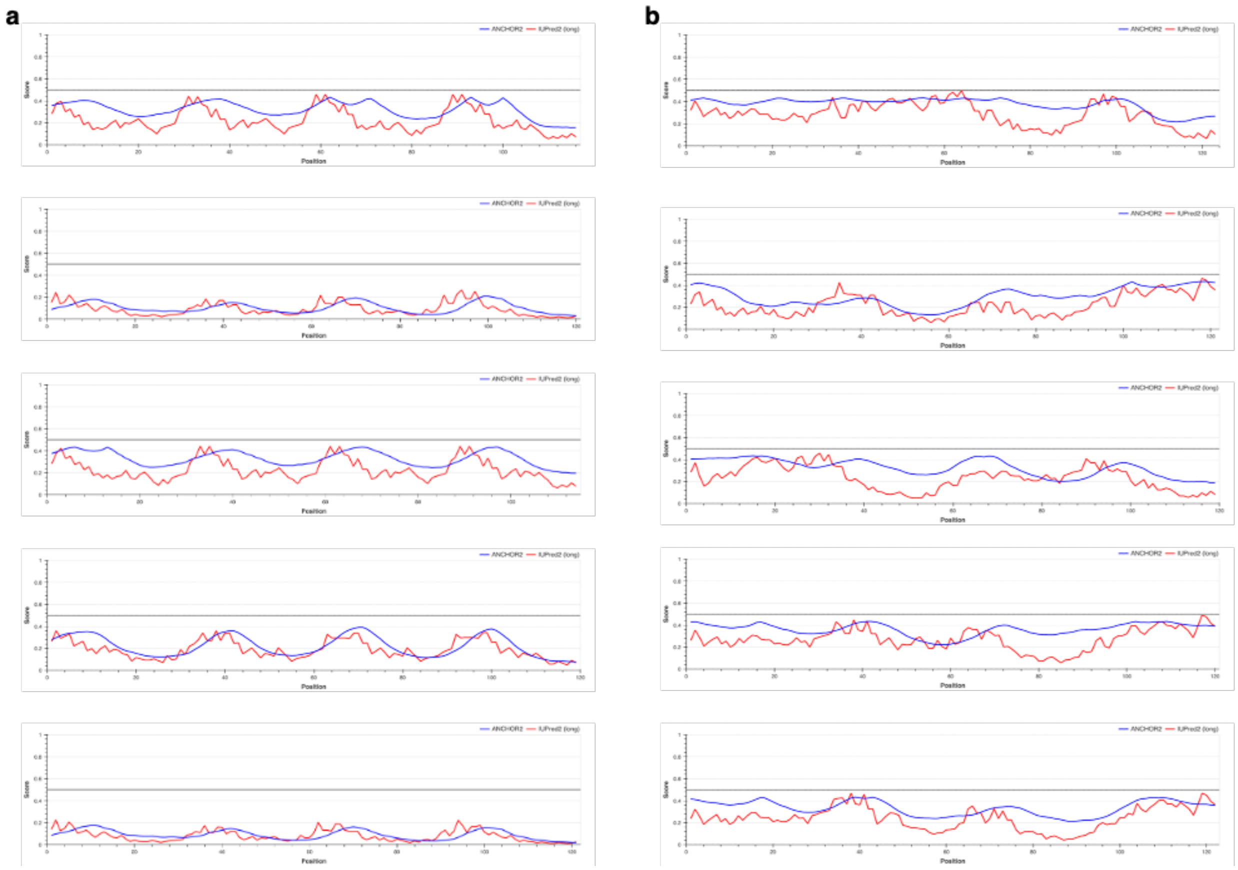


**Figure S8. Disorder measurements.** Comparison of (a) high self-similarity (cumulative percent identity: 560 - 600), to (b) low self-similarity (cumulative percent identity: 266 - 273). Highly self-similar clones show periodic disorder patterns with four evenly spaced peaks, closely resembling the TR-n4 profile. Protein disorder levels are measured using a web server. https://iupred2a.elte.hu
